# Excitatory neurons in stratum radiatum provide an alternative pathway for excitation flow that escapes perisomatic inhibition

**DOI:** 10.1101/2023.11.20.567832

**Authors:** Julia Lebedeva, David Jappy, Azat Nasretdinov, Alina Vazetdinova, Viktoria Krut’, Rostislav Sokolov, Yulia Dobryakova, Marina Eliava, Valery Grinevich, Andrei Rozov

**Author notes:** Correspondent author: Andrei Rozov Federal Center of Brain Research and Neurotechnologies, Moscow, Russia; OpenLab of Neurobiology, Kazan Federal University, Kazan, Russia; Department of Physiology and Pathophysiology, Heidelberg University, Heidelberg, Germany.

## Abstract

For over half a century, it has been postulated that the internal excitatory circuit in the hippocampus consists of three relay stations. Excitation arrives from the entorhinal cortex to the DG granule cells, is transmitted through the mossy fibers to CA3 pyramidal cells, and is then transmitted through Schaffer collaterals to CA1 pyramidal neurons. In all three structures (DG, CA3 and CA1), the activity of the excitatory neurons involved in the synaptic transmission of excitation are under the control of inhibitory basket neurons that are recruited into network activity via feed-forward and feed-back excitation. However, in the late 90s “stratum radiatum giant cells” were described as a novel type of neuron with the anatomical features of excitatory cells. Since then, the role of these cells in the hippocampal circuitry has not been well understood. Here, using optogenetic and electrophysiological techniques we characterized the functional location of these neurons within the hippocampal network. We show that: (i) the main excitatory drive to giant excitatory neurons in stratum radiatum (ExN_R_) comes via Schaffer collaterals; (ii) within the CA1 field, ExN_R_ are not directly connected with local pyramidal cells, but provide massive and efficient excitatory input to parvalbumin positive (PV+) interneurons; (iii) ExN_R_ are reciprocally innervated by bistratified cells, but not inhibited by backet interneurons; (iv) the efficiency of ExN_R_ excitation to PV+ interneurons is sufficient for a single ExN_R_ action potential to trigger massive inhibition of downstream CA1 pyramidal cells. Taken together, our data shows that ExN_R_ constitute an alternative pathway of excitation for CA1 interneurons that avoids the burden of perisomatic inhibition.

## Introduction

The hippocampus is one of the most studied structures in the CNS. The canonic trisynaptic hippocampal circuit appears in the famous drawings by Ramón y Cajal from the beginning of the 20^th^ century. Later, Andersen and Lomo proposed the chain of excitation flow that we know today: entorhinal cortex **→** *Dentate gyrus* granular cells **→** *Sratum pyramidale* CA3 pyramidal cells **→** *Sratum pyramidale* CA1 pyramidal neurons ^1^. Since then, hippocampal excitatory synapses have become the favored playground for long-term plasticity researchers. A simple PubMed cross search for “LTP” and “Hippocampus” results in more than 7500 papers ^2^. Even earlier, when working in Eccles laboratory, Per Andersen published a keystone paper describing perisomatic inhibitory neurons in the hippocampus and proposed the principles of recurrent inhibition ^3^. Later, basket cells were discovered and characterized for all three cellular populations constituting the hippocampal excitatory circuit ^4,5^. Merging the trisynaptic excitatory pathway with the feed-forward and feed-back inhibition provided by basket cells gave rise to multiple models of hippocampal rhythmic activity ^6–8^. Today, the generally accepted model of the functional-anatomical organization of excitatory circuits in the hippocampus assumes the same three main players that were drawn by Cajal and described in the pioneering work of Andersen, that is, DG granular cells and pyramidal neurons in the CA3 and CA1 regions. At the moment, in the search for diversification of hippocampal excitatory neurons, researchers seek unique anatomical, biochemical and molecular features that would allow subdivision of these three neuronal types into functionally distinct subpopulations. Recently, CA2 has emerged as an important area in the hippocampal circuitry, with distinct functions from those of CA3 ^9–11^. CA1 pyramidal neurons have been subdivided anatomically into “deep” and “superficial”, moreover, new findings show that the two subgroups are also functionally different ^12,13^. In the DG, as the site of adult neurogenesis, granule cells exhibit functional biochemical diversity depending on the stage of maturation ^14–16^. However, regardless of belonging to a particular subgroup, a common feature of neurons located in the pyramidal layer and DG is that they are under the inhibitory control of basket cells. In the basket cell population, fast spiking parvalbumin-positive basket cells were recognized as the main “dictators”, which completely determine rhythmogenesis ^17,18^ and also greatly diminish the efficiency of CA1 pyramidal cells as the main hippocampal output. A recent study by Hodapp et.al. suggested that neurons can escape the burden of perisomatic inhibition by relocating the axon initial segment from the heavily inhibited soma to one of the basal dendrites ^19,20^. While cells with a privileged dendrite possessing the axon can indeed bypass perisomatic inhibition, making one dendrite privileged makes the rest of dendritic tree obsolete. So, the question of who else can speak, when basket cells say: “Everyone be quite”, remains unanswered.

In 1996 Maccaferri & McBain described “giant cells” located in stratum radiatum that can express NMDAR-dependent LTP similarly to classical pyramidal cells ^21^. Two years later Gulyás and colleagues classified these neurons as excitatory cells based on a number of morphological features ^22^. Later it was shown that stratum radiatum excitatory neurons (ExN_R_) send their projections to the olfactory bulb ^23,24^. Thus, in addition to CA1 pyramidal neurons, the hippocampus has one more output neuronal population, and one more source of internal excitation, but in contrast to CA1 pyramidal cells, ExN_R_ are located in stratum radiatum, 100-300 um away from the innervation area of basket interneurons. Although the first description of ExN_R_ appeared more than 20 years ago, currently very little is known about the pre-and postsynaptic partners of these cells. Therefore, in this study, using an optogenetic approach and various electrophysiological techniques, we investigated the functional integration of ExN_R_ in the hippocampal circuitry.

## Results and Discussion

### Optogenetic identification the origin of excitatory synaptic inputs to ExN_R_

It has been shown that electrical stimulation in stratum radiatum can trigger EPSPs in ExN_R_ ^21^. However, this approach does not allow precise discrimination between Schaffer collateral inputs and entorhinal cortex projections. Especially, when taking into account that the apical dendrites of ExN_R_ extend to *stratum lacunosum moleculare* ^22^ (Fig 1A), which receives direct input from the pyramidal cells of layer III of the entorhinal cortex (EC) through the temporoammonal tract^25^. To overcome this limitation, we expressed hChR2-EYFP in either in CA3 or EC using in vivo AAV viral transduction. Two weeks after viral injection, when expression of hChR2 had reached a sufficient level, rats were sacrificed for slice preparation. ExN_R_ were recognized by the pyramidal-like appearance of the cell body, their location in *stratum radiatum* 100-300 μm away from the border of the superficial pyramidal cell layer and firing pattern (Fig 1A and B). In slices from rats with infected CA3 neurons, blue light pulses (1 ms), delivered to the area covering *stratum radiatum* and *stratum lacunosum molecul*are near the recoded cell, reliably triggered EPSCs in CA1 pyramidal neurons and ExN_R_ (Fig 1C). The latency of the responses relative to the onset of the light stimulus were nearly the same being 3.32±0.8 ms in ExN_R_ (n=8) and 3.26±0.5 ms in CA1 pyramidal cells (n=5). When hChR2 was expressed in EC neurons, EPSCs in ExN_R_ could be triggered in only 42% of the slices (n=12). Moreover, the averaged latency of the response was significantly longer (9.38±0.67 ms; n=5; p<0.001; Fig 1D and E) than that obtained when hChR2 was expressed in CA3 region. The low probability of getting responses upon stimulation of EC inputs together with the long polysynaptic latency suggest the absence of direct EC input to ExN_R_. Thus, CA3 pyramidal cells appear to be the main source of excitation for these neurons, at least within the hippocampal formation (Fig 1F).

**Figure 1.**
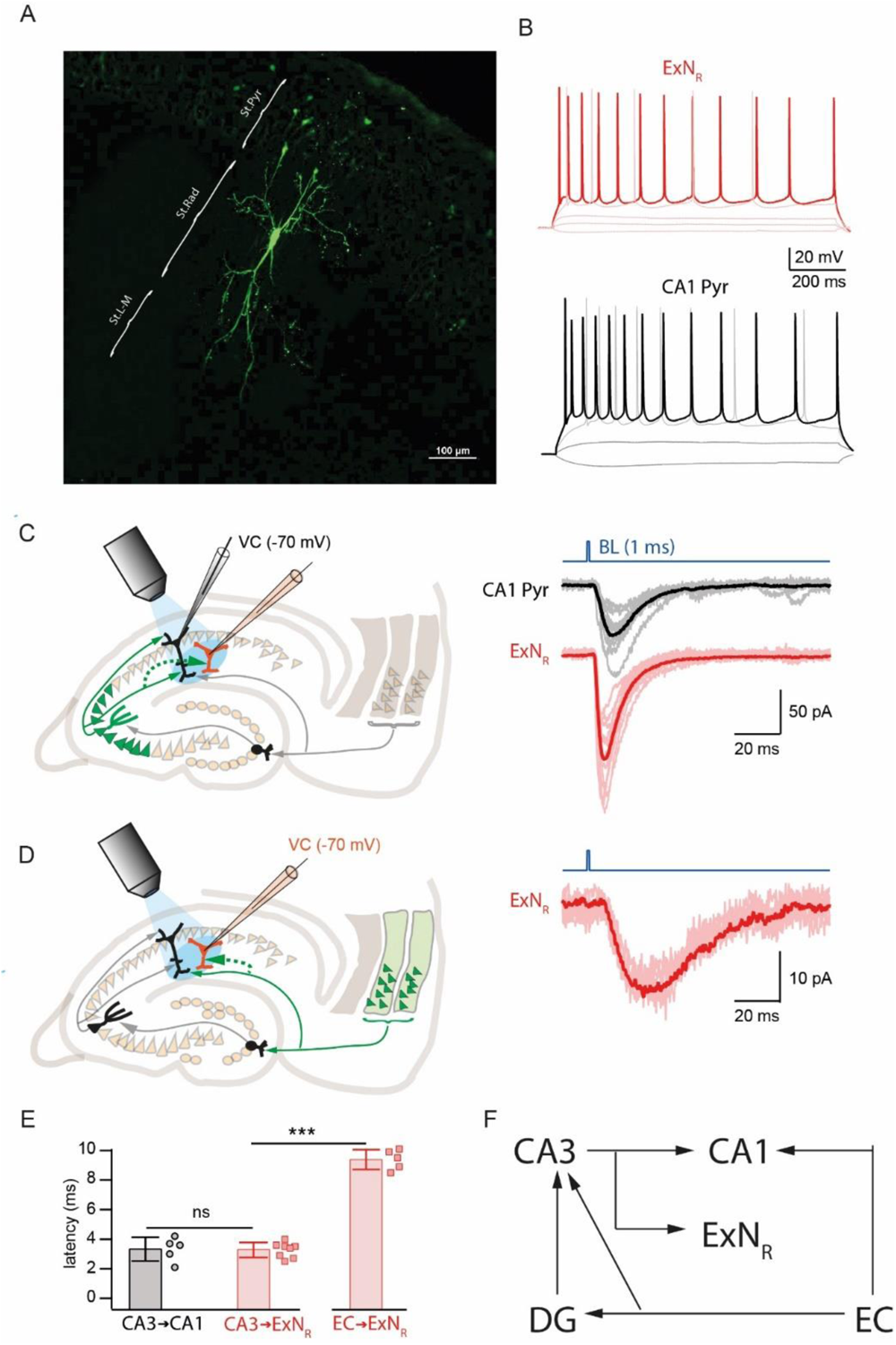
The place of ExN_R_ in the hippocampal excitation chain. A. z-Projected confocal images of ExN_R_ filled with Alexa 488. B. Typical voltage responses and firing patterns of ExN_R_ (red) and CA1 pyramidal neuron (black) triggered by hyperpolarizing and depolarizing current injections. C. Schematic drawing of experimental setup for optogenetic stimulation of CA3 pyramidal cell projections (left panel). Right panel shows blue light evoked individual and averaged responses in CA1Pyr (black) and ExN_R_ (red). D. Schematic drawing of experimental setup for optogenetic stimulation of EC projections (left panel). Example of blue light triggered individual and averaged IPSCs in ExN_R_ (right panel). E. Histogram compares latencies of EPSCs induced by optogenetic stimulation in CA1Pyr and ExN_R_. Averaged values ± SD presented by bars, symbols show data from individual experiments. F. Schematic diagram of excitatory inputs to major populations of hippocampal principal neurons and ExN_R_.

### Internal connectivity of ExN_R_ within CA1 circuitry

Next we explored the connectivity of ExN_R_with CA1 pyramidal neurons and two types of parvalbumin positive interneurons (Fig 2A): basket cells (BC) and bistratified neurons (Bist). We did not find direct connections between CA1 pyramidal cells and ExN_R_ in either direction (n=30). However, ExN_R_ innervated both types of interneurons with a relatively high connectivity rate (ExN_R_ to BC 36% and ExN_R_ to Bist 52%). Both types of excitatory synapses formed by ExN_R_ had very high initial release probability, rather large unitary EPSP amplitudes and showed prominent short-term depression (Fig 2A). For appropriate comparison we collected data on connectivity rate and synaptic efficacy characteristics for connections between CA1 pyramidal cells and the two types of interneurons under the same experimental conditions. Despite the fact that these pairs of pre-and postsynaptic neurons were located much closer to each other, both the chance of finding connected cells and the strength of connection were smaller when compared to those for ExN_R_ to BC and Bist connections (Fig. 2A and Fig. S2). Moreover, in 10% of connected ExN_R_ to Bist pairs and 7% of connected ExN_R_ to BC pairs, the amplitudes of evoked EPSPs could each the threshold level and trigger action potentials (AP) in the receiving postsynaptic cells. At connections formed by presynaptic CA pyramidal cells, the amplitudes of unitary EPSPs were substantially smaller and, therefore, the synchronized input from several pyramids is necessary for generation of a postsynaptic AP.

**Figure 2.**
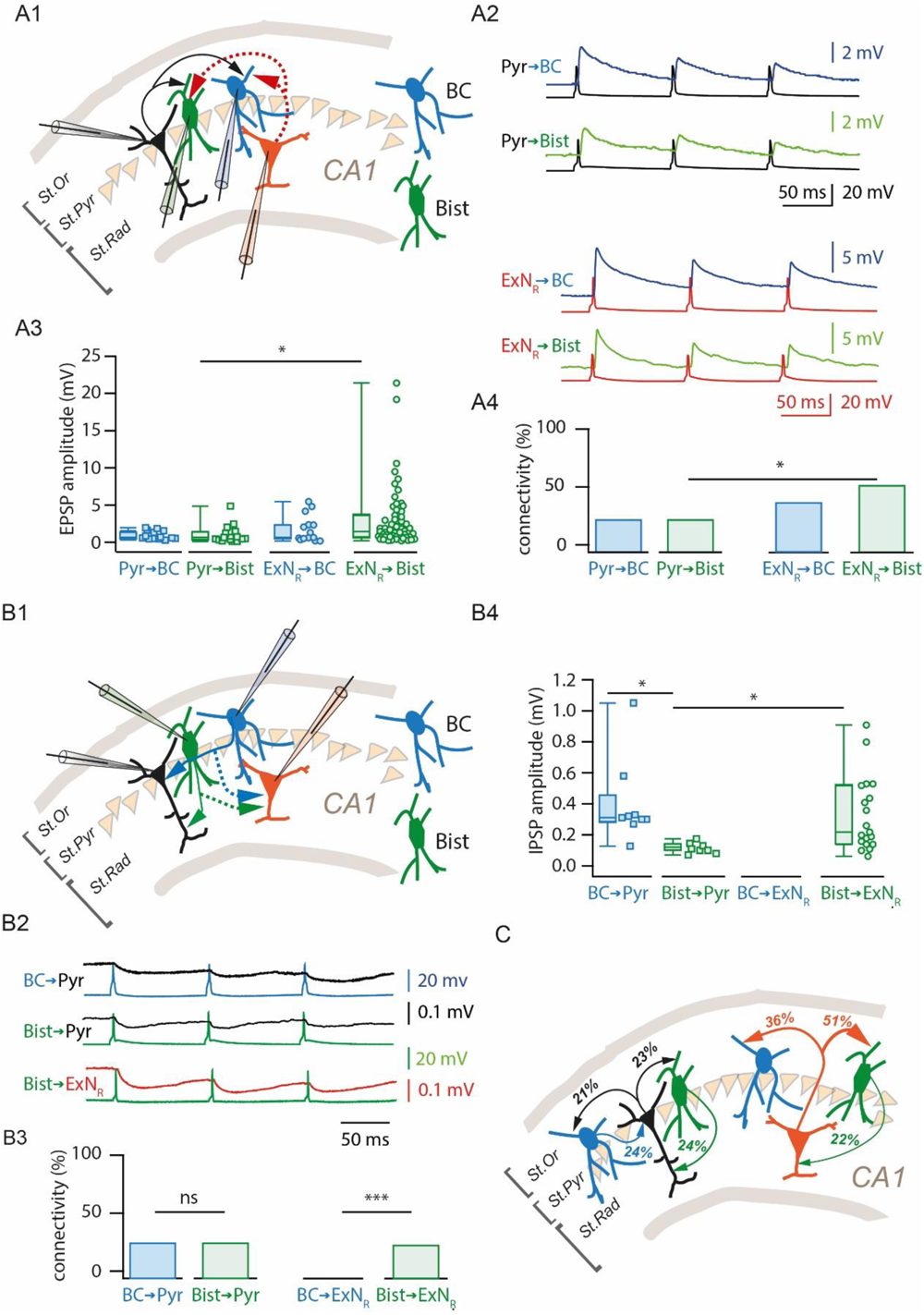
Synaptic communication of ExN_R_ with PV+ interneurons. A. Properties of excitatory projections from ExN_R_ and CA1Pyr to two types of PV+ interneurons. (A1) Diagram shows tested connections. (A2) Example averaged traces recorded from connected pairs between presynaptic CA1Pyr (Pyr) or ExN_R_ and postsynaptic basket cells (BC) or bistratified neurons (Bist). (A3) The plot shows median EPSP values together with the averaged EPSP amplitudes obtained in individual cell pairs formed by presynaptic CA1Pyr or ExN_R_ and postsynaptic BC or Bist. (A4) Comparison of connectivity rates at excitatory projections from CA1Pyr or ExN_R_ to two types of PV+ interneurons. B. Properties of inhibitory projections from *stratum oriens* PV+ interneurons to ExN_R_ and CA1Pyr. (B1) Diagram shows tested connections. (B2) Example averaged traces recorded from connected pairs between presynaptic PV+ interneurons and postsynaptic CA1Pyr (Pyr) or ExN_R_. Note the absence of connections from BC to ExN_R_. Comparison of connectivity rates (B3) and IPSP amplitudes (B4) for the inhibitory projections. C. Diagram summarizes data on connectivity within local circuitries formed by CA1Pyr and ExN_R_ with two types of stratum oriens PV+ interneurons. Note, while connectivity rates at reciprocal connections between CA1Pyr and BC or Bist are very similar, excitatory projections from ExN_R_ dominate over recurrent inhibition.

While searching for excitatory inputs from ExN_R_ and CA1 pyramidal cells to BC and Bist we also analyzed reciprocal GABAergic connections from these two types of interneurons (Fig. 2B). As expected, both BC and Bist innervate CA1 pyramidal cells with a connectivity rate of about 20-25%. The efficacy at perisomatic synapses was significantly higher than that at dendritic synapses formed by Bist (Fig 2B). However, the most intriguing question was whether ExN_R_ can be inhibited by BC. We didn’t find any connection out of 42 tested BC to ExN_R_ pairs, while connectivity from dendritic inhibitory Bist to ExN_R_ was almost as frequent (22%) as connections to CA1 pyramidal cells (25%). IPSP amplitudes were significantly higher at Bist synapses onto ExN_R_ as than those onto CA1 pyramidal neurons (Fig. 3B).

**Figure 3.**
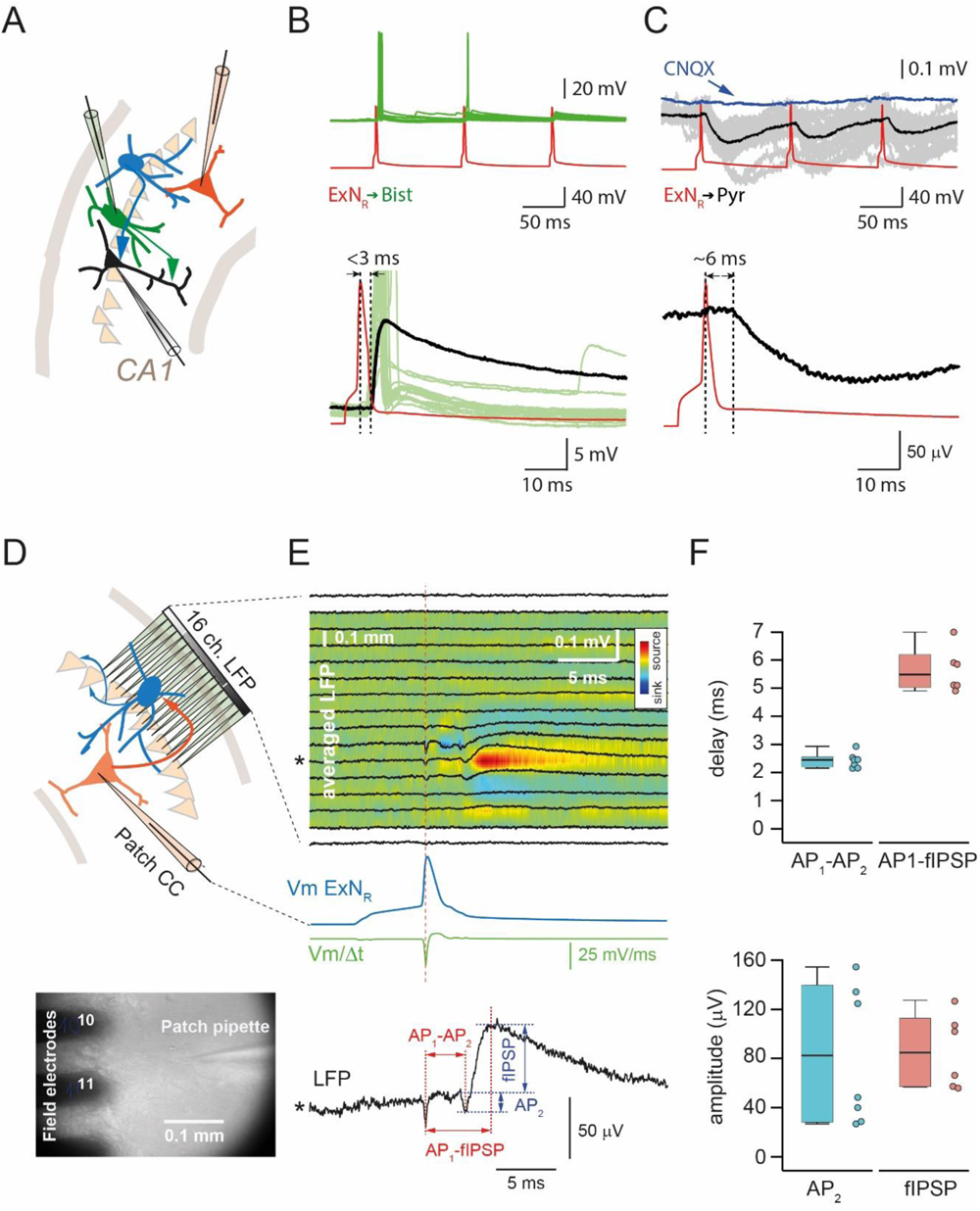
Excitatory drive from a single ExN_R_ is sufficient to trigger polysynaptic inhibition. A. Diagram shows connections tested for polysynaptic inhibition. B. Example of connected ExN_R_ **→** Bist pair where efficiency of synaptic transmission was sufficient to reliably trigger a postsynaptic AP. Subthreshold EPSPs were observed in 10% of ExN_R_ **→** Bist and 7% 0f ExN_R_ **→** BC pairs. C. Polysynaptic IPSPs recorded at ExN_R_ **→** CA1Pyr. Recruitment of an extra interneuron that receives excitatory drive from the stimulated ExN_R_ and innervates the recorded CA1Pyr was confirmed by the disynaptic delay of IPSPs relative to the peak of AP and the complete occlusion of IPSPs in the presence of an AMPAR antagonist (CNQX; blue trace). D. Schematic representation of the experimental setup (upper panel): 16-channel electrode array for local field potential (LFP) recording from the pyramidal layer and patch-clamp recording from the ExN_R_ in striatum radiatum. The micrograph (bottom panel) of the hippocampal slice shows the location of the extracellular electrodes and patch pipette. E. Averaged LFP and current source density profile in the hippocampal pyramidal layer relative to the peak of evoked action potential (AP) of the ExN_R_ (upper panel). Averaged AP of the ExN_R_ (bottom panel; blue trace), LFP for channel #11 (black trace, also marked with an asterisk in the top panel), and the first derivative of Vm of the averaged AP waveform (Vm/Δt, green trace). The LFP trace shows the 4 parameters calculated for all experiments (AP_1_ to AP_2_ delay, AP_2_ amplitude, fIPSP amplitude, AP_1_ to fIPSP peak time) AP_1_ is the negative peak on the LFP that coincides in shape with the first derivative of the AP (Vm/Δt) and therefore represents LFP fluctuation caused by evoked AP in ExN_R_. AP_2_, following AP_1_ with a characteristic monosynaptic delay, reflects activity in the second neuron excited by the stimulated ExN_R_. Finally, the fIPSP following AP_2_ suggests the GABAergic nature of the second neuron. (F) Box plots show medians (P_25_; P_75_) and corresponding individual values for the following parameters: AP_1_ to AP_2_ delay and AP1 to fIPSP delay (upper panel); AP_2_ and fIPSP amplitudes, (bottom panel).

These data allow ExN_R_ to be considered “privileged neurons” that escape perisomatic suppression by BC while providing massive excitatory drive to both BC and Bist, thus promoting inhibition of CA1 pyramidal cells (Fig. 2C).

### ExN_R_ can trigger feed-forward inhibition of CA1 pyramidal cell layer

As mentioned above, a single AP in a presynaptic ExN_R_ can trigger APs in the postsynaptic BC and Bist (Fig 3B). Although, we have not found any evidence of direct synaptic input from ExN_R_ to CA1Pyr, in 10% of the tested pairs stimulation of ExN_R_ could trigger a disynaptically delayed IPSP (delay: 6.4±0.4 ms; amplitude: 0.08±0.24 mV; n=3; Fig 3C) in CA1Pyr. Application of 10 μM of CNQX, an AMPAR antagonist completely blocked IPSPs, suggesting that stimulation of ExN_R_ could recruit interneurons projecting to CA1Pyr. Thus, ExN_R_ can operate as an amplification relay station for feed-forward inhibition of neurons in the CA area.

To examine this notion, we tested the effect of single ExN_R_ stimulation on the local field potential profile across the CA1 pyramidal layer recorded using a 16 channel multishank silicone probe (Fig 3D). In 24% of the slices unitary APs in ExN_R_ generated an fIPSP, delayed relative to peak of the AP by 5.5 ms (n=6; Fig 3D-F). Moreover, often fIPSPs were preceded by a secondary spike with a delay characteristic for monosynaptic transmission (2.6 ms; n=7; Fig 3D-F).

### Possible functions of ExN_R_ within hippocampal network

Although, previous studies suggested that ExN_R_ are excitatory neurons, this assumption was made based on morphological features and the absence of histological markers typical for interneurons ^22,23^. Here we directly show the glutamatergic nature of ExN_R_ by recording from pairs of connected cells. It is generally accepted that AP generation in CNS requires either high frequency stimulation or unitary input integration summation of multiple inputs^26^. Low effectiveness at individual excitatory connections was suggested to be essential for temporal and spatial information coding and, therefore, for increased computational capacity of neuronal networks. We found that ExN_R_ often do not follow this rule at their synapses converging on BC and Bist neurons. It has to be emphasized here, that all experiments were done on brain slices where truncation of axons during slicing affects both the chances of finding connections and synaptic efficacy. Obviously, the impact of negative artefacts of slicing increases as a function of distance between tested neurons. Thus, the efficiency of ExN_R_-driven excitation should be substantially higher in the intact hippocampal network. Another unique trait of ExN_R_ is that in contrast to the other hippocampal excitatory cells, they are not controlled by *stratum oriens* BCs (Fig 4A). Thus, these two features suggest ExN_R_ play an exclusive role within the hippocampal circuitry.

**Figure 4.**
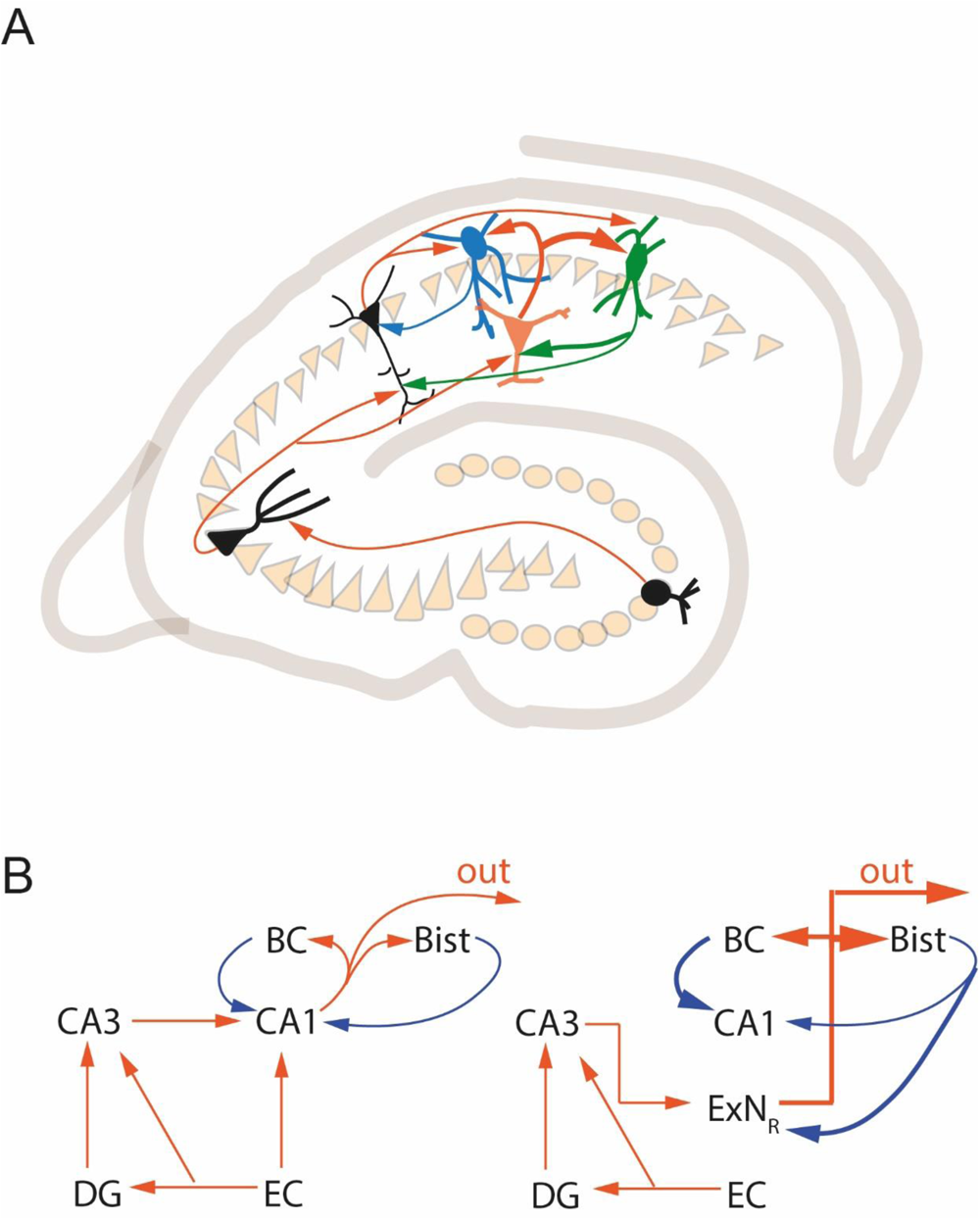
Possible functional role of ExN_R_ in control of perisomatic inhibition of hippocampal output. A. Schematic drawing depicting excitatory and inhibitory inputs of ExN_R_ and their synaptic targets. The main differences from CA1Pyr are (i) the absence of external excitatory input from EC and (ii) a lack of inhibitory control provided by PV+ backet cells. B. The left diagram shows the classical view of excitatory and inhibitory circuitries that control the major hippocampal output (CA1 pyramidal cells). Excitation converges from EC via the trisynaptic pathway and directly through the temporoammonal tract and is efficiently controlled by means of feed-back inhibition, primarily via reciprocal connections with BC. The balanced interplay between excitation and inhibition of pyramidal cells was postulated to be the main structural prerequisite for hippocampal rhythmogenesis. However recruitment of ExN_R_ (right diagram), bypassing BC-mediated perisomatic inhibition, can shift the excitation/inhibition balance of CA1Pyr and disrupt the functioning of the CA1Pyr-BC oscillator.

Reciprocal connections between excitatory cells located in *stratum pyramidale* and BC form a chain of micro-oscillators that can generate, maintain and transmit both normal and pathological rhythmic activities. One can assume that ExN_R_, being not phase-locked by BC-driven synchronization, can disturb the oscillatory activity by activating interneurons out of phase. This might be an important mechanism to prevent epileptiform activity. The second scenario of possible ExN_R_ network function comes from the fact that both ExN_R_ and a subpopulation of CA1Pyr project to the olfactory bulb. Thus, ExN_R_ especially located in the ventral aspect of the hippocampus may emphasize their own input to the olfactory bulb by simultaneous inhibitions of CA1Pyr also projecting to this region (Fig 4B).

The exact function of ExN_R_ remains to be found, but the existence of one more powerful source of excitation that can bypass perisomatic inhibition should be taken in account in hippocampal network models.

## Methods

### Preparation of rat brain slices

Horizontal brain slices (400 µm thick) containing the hippocampus and entorhinal cortex were obtained from male Wistar rats 6–8 weeks of age using a standard procedure^27^. All experimental protocols were approved by the State Government of Baden-Württemberg (Projects T100/15 and G188/15) or by the Local Ethical Committee of Kazan Federal University (#24/22.09.2020). Rats were killed under deep CO_2_-induced anesthesia. After decapitation, brains were rapidly removed and placed in cold (1–4°C) oxygenated artificial CSF (ACSF) containing the following (in mM): 124 NaCl, 3 KCl, 2 CaCl_2_, 1 MgCl_2_, 10 glucose, 1.25 NaH_2_PO_4_, and 26 NaHCO_3_, saturated with carbogen (95% O_2_ and 5% CO_2_), with pH 7.4 at 34°C. Horizontal brain slices containing the intermediate/ventral portion of the hippocampus and connected areas of the entorhinal cortex were cut using a vibratome slicer (VT1200S, Leica). In optogenetic experiments for better preservation of connectivity between the entorhinal cortex and hippocampus, slices were cut with at an angle of ∼15° toward the ventral side. Before electrophysiological recordings, slices were allowed to recover for at least 1 h. Slices were kept in a submerged incubation chamber at room temperature.

### Injection of virus and optogenetic stimulation

All experiments requiring injections of AAV were conducted in a biosafety level 1 laboratory. The AAV5-CaMKIIa-hChR2(H134R)-EYFP was injected bilaterally into CA3 and the entorhinal cortex at 1 μl/site. During operations, rats were deeply anesthetized with isoflurane (4%) and mounted in a stereotaxic frame. Anesthesia was maintained by mask inhalation of vaporized isoflurane at concentrations between 1.5% and 2.5%. Following head fixation, the skull was exposed and a small burr hole was drilled above the injection site. The AAV was infused using amicroinfusion pump (Stoelting Co., USA) at a rate of 0.2 µl/min. The bilateral injections were performed stereotaxically into the CA3 area (5.2 mm posterior, 4.4 mm lateral to bregma, 7.0 mm dorsoventral) and into the entorhinal cortex (10.0 mm posterior, 4.6 mm lateral to bregma, 8 mm dorsoventral, the needle was inserted at the angle of 16-20°) (Paxinos and Watson, 1998) using a 10 μl Hamilton syringe (Hamilton, USA). The first point of injection into EC was at 8.0 mm dorsoventral (0.5 ml), then the needle was retracted to 5.0 mm and an additional 0.5 ml of virus was injected. The needle was left in place for another 10 min before it was withdrawn. When all injections were completed, the wound was sutured and the animal was monitored during recovery from anesthesia, after which it was returned to its home cage. Animals were allowed to recover for a minimum of 2 weeks after injections, before being sacrificed.

The axonal fibers expressing ChR2 were excited through a 40×/0.8 numerical aperture (NA) objective using a transistor–transistor logic-controlled blue LED (470 nm; pulses, 1ms; catalog #M470L3, ThorLabs). Neurons were recorded in VC mode and held at −70 mV using a K gluconate-based pipette solution containing 144 K-gluconate, 4 KCl, 10 HEPES, 4 Mg-ATP, 0.3 Na-GTP, and 10 Na_2_-phosphocreatine, adjusted to pH 7.3 with KOH. GABAergic synaptic transmission was blocked by the continuous presence of the GABA_A_ receptor antagonist SR95531 (10 μM).

### Synaptic connectivity of ExN_R_

ExN_R_ and CA1Pyr were identified by firing pattern and location in the slice, in stratum radiatum and stratum pyramidale respectively. BC and Bist interneurons were distinguished by firing pattern (Fig S1) and input resistance (IR). BC had significantly lower IR (Median: 103 MOhm; n=30) and required at least 150-200 pA of current injection for spike generation. IR in Bist interneurons was 191 MOhm (n=30; p<0.001) and 50 pA of current injection could trigger AP firing. To substantiate our discrimination criteria 5 neurons of each group were filed with biocytin, and the identity of interneurons was confirmed by axonal arborization pattern. Dual whole-cell recordings were performed at room temperature. Slices were continuously superfused with an extracellular solution containing the following (in mM): 125 NaCl, 2.5 KCl, 25 glucose, 25 NaHCO_3_, 1.25 NaH_2_PO_4_, 2 CaCl_2_, and 1 MgCl_2_, bubbled with 95% O_2_/5% CO_2_. To study synaptic connections, presynaptic cells were stimulated with a 10 Hz train of three suprathreshold current pulses, which were repeated every 7 s. All paired recordings used for connectivity analysis were conducted in CC mode. During recordings, cells were held at resting membrane potential. Averages of 50–100 consecutive sweeps were used for the analysis of postsynaptic responses.

### Combined LFP and whole cell recording

Experiments with multichannel LFP recordings combined with single ExN_R_ subthreshold stimulation were done at 33°C. A 16 channel multishank silicone probe was positioned in the CA1 pyramidal cell layer along stratum pyramidale.

Signals were recorded using an Open Ephys Acquisition Board (Cambridge, Massachusetts) at a sampling rate of 30 kHz. For stimulation we chose ExN_R_ that were located in front of the center of the multishank probe, usually between the 8^th^ and 11^th^ electrodes, and at least 100 μm away from the pyramidal cell layer. The cells were repeatedly stimulated by depolarizing pulses, sufficient to trigger a single AP. Interpulse interval was 7 seconds. Total number of stimuli in every experiment was 50-100.

To avoid erroneous detections, the ExN_R_ action potential waveforms were detected as local peaks of the second derivative of Vm, then the maximum value of Vm was determined in the 8 ms window after each Vm’’ peak. The corresponding AP peak time was used to construct an averaged LFP and current source density profile in the [-10 to 20] ms window. Four parameters were calculated from the averaged LFPs: AP_1_ to AP_2_ peak to peak delay time, AP_2_ amplitude (second negative peak), fIPSP amplitude (maximum positive peak), AP_1_ to fIPSP peak to peak dealay time, where AP_1_ is the first negative peak on LFP that coincides in shape with the first derivative of the ExN_R_ AP (Vm’).

### Statistical analysis

Data in figures are presented as medians (P_25_; P_75_) and individual values, unless otherwise stated. Whiskers show minimum and maximum values. Statistical analysis was performed using SigmaPlot (Systat), GraphPad (InStat, GraphPad Software) or Matlab Statistics Toolbox. Mann–Whitney *U test*, Wilcoxon Rank Sum test or Fisher’s exact test were used for statistical comparisons A *p* value <0.05 was regarded as significant (for all data: \**p* < 0.05, \*\*\**p* < 0.001, ns, not significant).

## Acknowledgements

We thank Dr. Andreas Draguhn and Dr. Oliver Kann for infrastructural support of this project and Irina Kopylova for the invaluable help with image processing. This work was supported by “Center of Photonics” funded by the Ministry of Science and Higher Education of the Russian Federation (contract no. 075-15-2022-293) to DJ, RS and AR

## Supplementary figures

**Figure S1.**
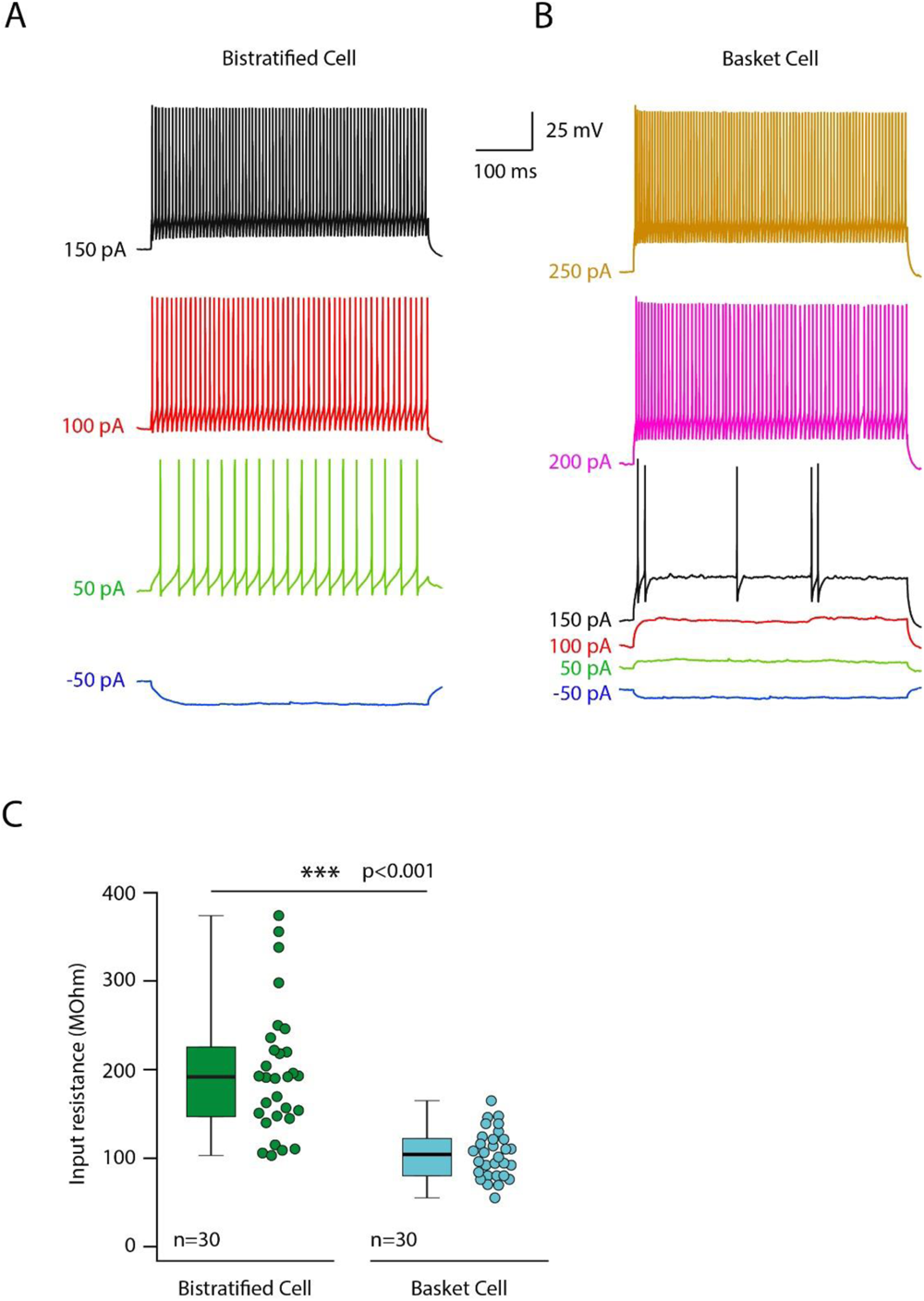
Intrinsic properties of bistratified and basket cells. A. Voltage responses and spiking to hyperpolarizing and depolarizing 1 second current injections in typical bistratified neuron. Note that minimal amplitude (50 pA) of injected current is sufficient to trigger sustained firing throughout whole period of depolarization. B. Voltage responses and spiking to hyperpolarizing and depolarizing 1 second current injections in typical basket cell. Note that much higher amplitude (200 pA) of injected current is needed to trigger sustained firing throughout whole period of depolarization. C. Box plot compares medians (P_25_; P_75_) and corresponding individual input resistance values in bistratified and basket cells.

**Figure S2.**
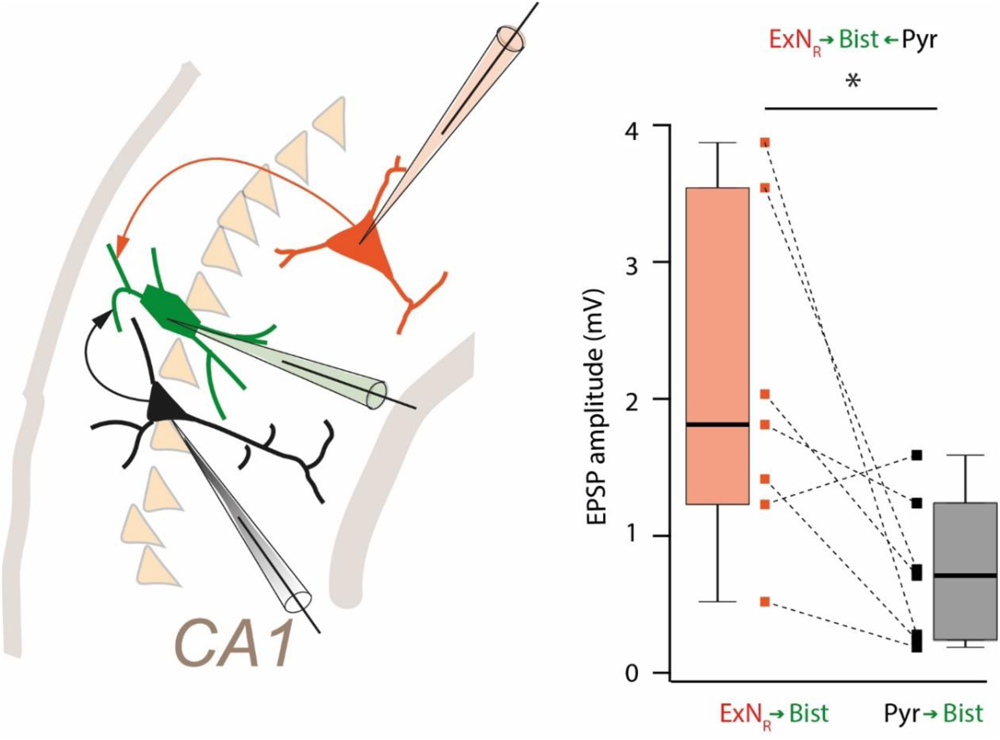
Direct comparison of synaptic efficacy at ExN_R_ → Bist and CA1Pyr → Bist synapses. Diagram shows experimental design of sequential triple recordings. After finding synoptically connected pair from either ExN_R_ or CA1Pyr to Bist, we searched for the alternative excitatory input (CA1Pyr or ExN_R_, respectevly) to the same interneuron. Box plot provides pairwise comparison of the avareged EPSP amplitude at two types of excitatory synapses (from presynaptic ExN_R_ and CA1Pyr) converging on the same postsynaptic bistratified cell.

